# Quantification and statistical comparison of cell-state transition kinetics using a parametric failure-based model

**DOI:** 10.64898/2026.04.21.719724

**Authors:** Stanley E. Strawbridge, Alexander G. Fletcher

## Abstract

Successful development of multicellular organisms requires cells to transition between states with precise timing. Distinct cell states are often understood as being maintained by stabilizing regulatory networks, such that a complete cell-state transition requires network rewiring through partial dismantling of the current state and concurrent reconfiguration into a new one. Empirically, these transitions are often investigated by quantifying the gain or loss of expression of a small number of state-specific markers, frequently a single proxy. A general quantitative framework for comparing the kinetics of such transitions across experimental conditions is lacking. Here, we show that the delayed Weibull distribution provides a natural description of cell-state transition kinetics when transition is viewed as the cumulative consequence of many molecular events, whose timing may vary between cells and conditions, analogous to system failure in reliability theory. This formulation yields a compact model with interpretable parameters describing the delay before transition onset, the characteristic timescale of transition, the temporal form of the transition hazard, and the fraction of cells competent to respond. Together, this framework provides a practical and interpretable approach for quantifying the kinetics of cell-state transitions and how they are altered by perturbation.

## INTRODUCTION

The two defining features of stem cells are self-renewal - their ability to continuously generate at least one unaltered daughter cell of equivalent potency - and differentiation - the capacity to give rise to progeny with more restricted potential (Smith, 2006). Stem cells maintain their state indefinitely when supported by specific combinations of pro-self-renewal signals and anti-differentiation signals. *In vivo*, this is facilitated by the stem cell niche, exemplified by hematopoietic stem cells in the bone marrow (Morrison and Scadden, 2014), intestinal stem cells in the crypt (Clevers, 2013), neural stem cells in the subventricular zone (Chaker et al., 2024), and spermatogonial stem cells in the testis (Bhang et al., 2018).

Similarly, certain stem cells can be maintained *in vitro* through exogenous growth factors and small-molecule inhibitors that stabilise a specific cellular identity. Pluripotent stem cells provide a primary example; distinct, stable pluripotent states have been captured *in vitro* across various species. These states correspond to different developmental stages (Nichols and Smith, 2009) and are obtained via direct derivation or the reprogramming of cells with more restricted potential (Evans and Kaufman, 1981; Martin, 1981; Thomson et al., 1998; Brons et al., 2007; Tesar et al., 2007; Takashima et al., 2014; Guo et al., 2016; Takahashi and Yamanaka, 2006). Such states are maintained by a regulatory network of interacting components - including transcription factors, chromatin states, and signalling pathways - that collectively sustain functional identity. Upon the withdrawal of self-renewal signals and the introduction of pro-differentiation cues, this regulatory network undergoes reconfiguration. We define a cell-state transition as the point of entry into a network configuration where previous functions are irreversibly lost and the acquisition of new functions are initiated.

As these regulatory networks comprise many interacting molecular components, the exit from a stable cell state is unlikely to be a single deterministic event. Instead, transition is a consequence of the progressive destabilisation of the regulatory network, driven by the accumulation of molecular changes that erode the interactions sustaining the original identity. Because the timing of this destabilisation varies between cells - even within seemingly homogeneous populations - transitions are more accurately described by a distribution of transition times rather than a single characteristic value (Lebek et al., 2024; Strawbridge et al., 2026). At the population level, this distributed timing manifests as a gradual change in the proportion of cells that have completed the transition. This suggests that cell-state transition kinetics may be described using probabilistic frameworks developed for systems in which macroscopic change emerges from the accumulation of numerous microscopic events.

Previous mathematical treatments of cell-state transitions have sought to formalise the shift from discrete cell-type classifications to the continuous, high-dimensional trajectories observed in single-cell data. As highlighted by Mulas et al. (2021), a key challenge is defining transition boundaries when cell states are viewed as metastable configurations within a dynamic continuum. To address this, ‘landscape’ models have gained prominence, using statistically derived geometric potentials to capture the logic of cellular decision-making dynamics (Sáez et al., 2022). In these frameworks, cell-state transitions are represented as movements across a manifold, where topographical features - ridges and valleys - govern the probability and direction of transitions. Complementing these geometric approaches, stochastic branching models have been used to disentangle the coupling between differentiation and cell division. For example, Ruske et al. (2020) showed that transition rates and division kinetics can be inferred from clonal snapshots, enabling quantification of how proliferation influences the proportion of differentiated progeny.

However, existing models tend to focus on the logic of fate decisions or mean population-level rates, rather than the explicit temporal mechanics of cell-state transitions. They often assume Markovian dynamics with constant transition probabilities or rely on high-dimensional fits that are difficult to interpret in terms of the progressive degradation of regulatory networks. This raises a key question: how can we quantify the stochastic timing of cell-state transitions in a way that reflects the cumulative destabilisation of the underlying molecular architecture?

Here, we address this gap by applying the delayed Weibull statistical distribution (Weibull, 1951) to quantify cell-state transition kinetics. Drawing a formal analogy to reliability engineering, we model cell-state transitions as the cumulative result of probabilistic molecular changes that destabilise the regulatory interactions maintaining the initial state, a process comparable to system failure in complex engineering structures. In this sense, cell-state transition is treated as an effective failure event. When assays measure loss of the initial state directly, this interpretation is straightforward. When assays instead measure acquisition of the next state, the model assumes that productive transition requires both collapse of the initial state and establishment of the new one. Accordingly, the fitted kinetics should be interpreted as describing effective completion of the rewiring process at the level of the measured phenotype, rather than decay of the initial network alone. This distinction is important because exit from the first state is not by itself sufficient for successful transition, and cells may instead abort if the post-transition network is not installed (Mulas et al., 2024).

We present a parametric model, derived from this failure-based analogy, to describe the temporal progression of cell-state transitions. The resulting delayed Weibull formulation provides a compact functional form with interpretable parameters describing the onset delay, the characteristic timescale of transition, the temporal form of the transition hazard, and the fraction of cells competent to respond within the observation window. We apply this framework to published datasets spanning marker loss during differentiation, lineage acquisition, and cellular reprogramming, and use it to compare transition kinetics across experimental conditions. Ultimately, this approach provides a practical tool for quantifying the kinetics of cell-state transitions and analysing how they are altered by perturbations.

## RESULTS

### A delayed Weibull distribution describes key aspects of cell-state transition kinetics

We model a cell state as maintained by a stabilising regulatory network of interacting molecular components. Each component can fail to support the current state through stochastic events (Fig. 1A; see also Supplemental Model, Section 2.1). In this framework, a cell-state transition is the cumulative result of multiple failure events, rather than the result of a single deterministic step. A successful transition requires both the collapse of the original state and the successful installation of the new one. This perspective is motivated by our previous analysis of exit from naïve pluripotency, where a time-dependent failure process accurately described state-transition kinetics (Strawbridge et al., 2026). To obtain a tractable population-level description, we make three primary assumptions: (i) individual failure events are conditionally independent given the state of the system; (ii) at the population level, failure contributions to network destabilisation accumulate additively; (iii) following a common onset delay, each node is shares the same general temporal form of failure, differing only in scale. Under these assumptions, the probability that a cell has transitioned by time *t* follows a delayed Weibull distribution (Weibull, 1951). To account for cells that may not be competent to transition within the observation window, we scale this distribution by a competence parameter *π*, so that the probability that transition occurs before time *t* is given by

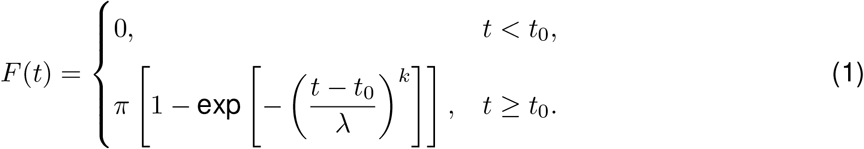

**Figure 1.**
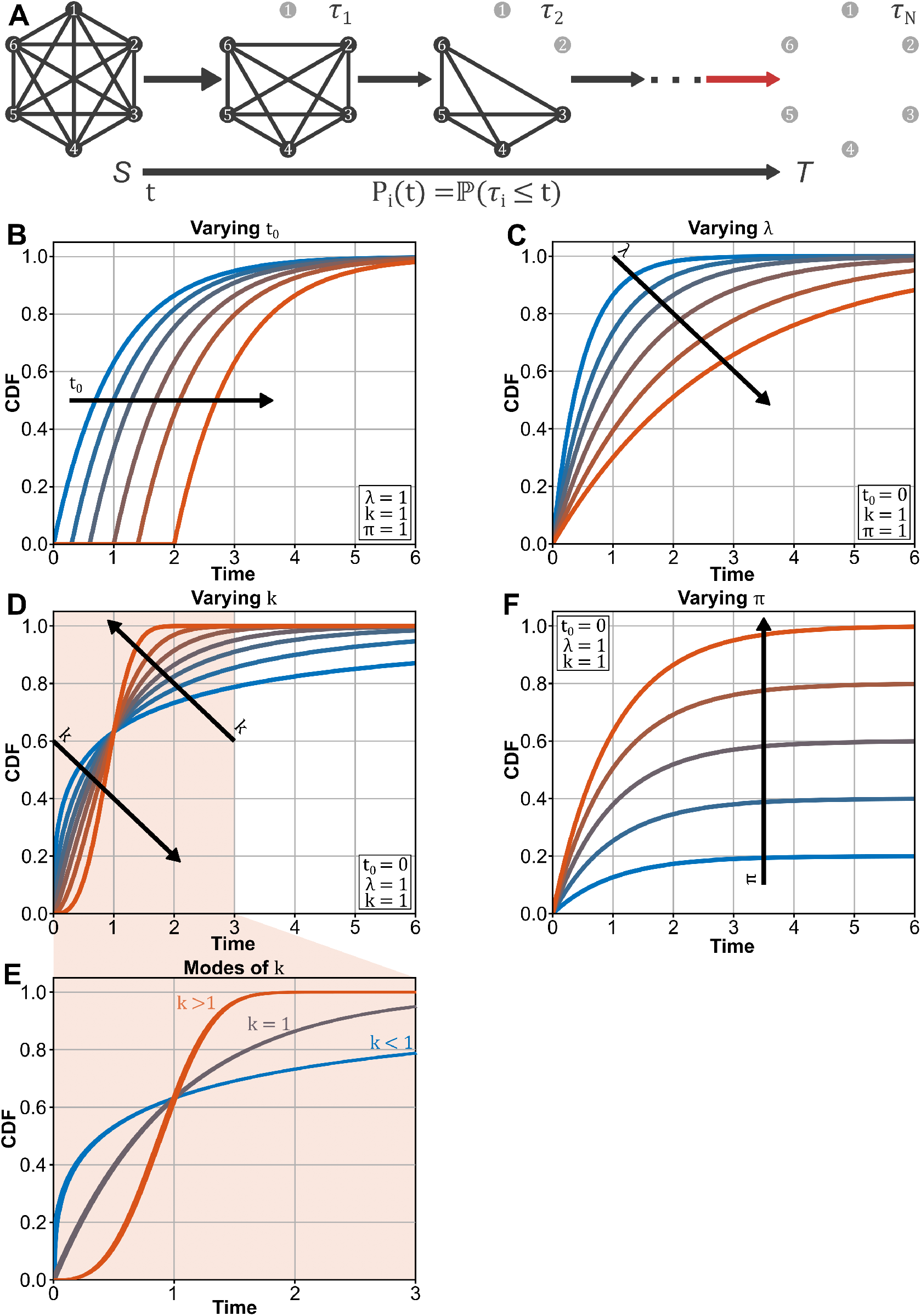
A delayed Weibull framework separates cell-state transition kinetics into interpretable features of onset, pace, hazard, and response extent. **A**. Schematic of a regulatory network that stabilises a cell state. Nodes represent molecular components whose collective interactions maintain cellular identity. Progressive node failure destabilises the network and permits transition to an alternative state. **B**. Increasing *t*_0_ delays the onset of transition without changing the overall shape of the transition profile. **C**. Increasing *λ* slows accumulation of the transitioned fraction and therefore delays the overall pace of transition. **D**. The shape parameter *k* determines the temporal form of the hazard, allowing decreasing, constant, or increasing transition propensity over time. **E**. The three regimes *k <* 1, *k* = 1, and *k >* 1 correspond to decreasing, constant, and increasing hazard, respectively, and generate qualitatively distinct transition profiles. **F**. Increasing *π* raises the asymptotic transitioned fraction without altering the temporal form of the profile among competent cells.

This formulation decomposes cell-state transition kinetics into four distinct, interpretable features (see also Supplemental Model, Section 2.2): *t*_0_ specifies the latency period before measurable state change begins (Fig. 1 B); *λ* sets the characteristic timescale, with larger values indicating slower progression (Fig. 1 C); *k* determines how transition propensity changes over time (Fig. 1 D); and *π* controls the fraction of cells able to respond (Fig. 1 F).

The hsape parameter *k* is particularly informative, as it admits three modes corresponding to different temporal organizations (Fig. 1 E): *k* = 1 corresponds to a memoryless process with constant hazard rate; *k >* 1 to an increasing hazard, where transition becomes progressively more likely; and *k <* 1 to decreasing hazard, indicating early susceptibility followed by a slowing rate.

This parameterization yields experimentally useful summary statistics (see Supplemental Model, Section 2.3). The delay *t*_0_ enters additively into characteristic transition times, shifting the process without altering its shape. The scale parameter *λ* sets the overall pace, scaling both the mean and variance of the competent subpopulation. While *π* determines the ultimate responding fraction, it does not affect the timing of transitions among those competent cells. Since the delayed Weibull distribution is available in closed form, characteristic transition times - such as *t*_10_, *t*_50_, or *t*_90_ - can be calculated analytically. In practice, *t*_50_ (the time at which 50% of the competent population has exited the initial state) is often more intuitive to interpret and compare across experimental conditions than *λ* itself. Together, these four parameters provide a flexible framework for quantifying when transitions begin, their pace, their temporal dynamics, and the total cellular response.

### Distinct modes of the shape parameter *k* arise from simple classes of network topology

The shape parameter *k* determines the time dependence of the hazard function for competent cells (see also Supplemental Model, Section 2.2):

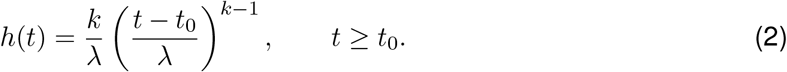

Accordingly, *k* determines whether the instantaneous probability of transition increases, decreases, or remains constant over time following the delay period.

To identify which network topologies generate distinct modes of *k*, we constructed minimal stochastic models for constant-, increasing-, and decreasing-hazard regimes that emulate different biological processes. We simulated a large ensemble of single-cell trajectories (10^8^ cells) for each model using the stochastic simulation algorithm (Gillespie, 1977). This scale was chosen to reflect the large effective cell numbers typically probed in pooled differentiation assays and to obtain smooth empirical cumulative distribution and hazard estimates for comparison with delayed Weibull fits. In all simulations, we set *t*_0_ = 0 and *π* = 1 to ensure that differences between models reflected the temporal form of the hazard rather than variations in onset delay or the competent fraction. In each simulation, a cell was initialised in the starting state and allowed to evolve until it first reached the transitioned state, *T*. These first-passage times were used to construct a cumulative distribution function for each topology, which was then fitted with the delayed Weibull model, mimicking the analysis of experimental population-level data.

When *k* = 1, the hazard is constant:

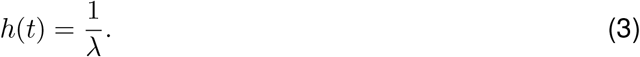

The instantaneous probability of transition is therefore independent of how long a cell has remained in the pre-transition state after *t*_0_. A minimal topology consistent with this regime is the one-step process

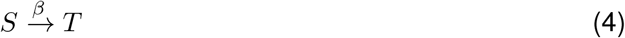

(Fig. 2 A). Stochastic simulation of this topology recovers a delayed Weibull fit with *λ* = 0.999832 and *k* = 1.000049. The resulting cumulative distribution function is exponential, as expected for a memoryless one-step process (Fig. 2 D, blue lines). This, in turn, gives rise to a hazard that is constant through time (Fig. 2 E, blue lines). Such a regime is expected when escape from the initial state *S* is controlled by a single effective rate-limiting event, or when the aggregate effect of many weakly interacting processes is approximately Poissonian.

**Figure 2.**
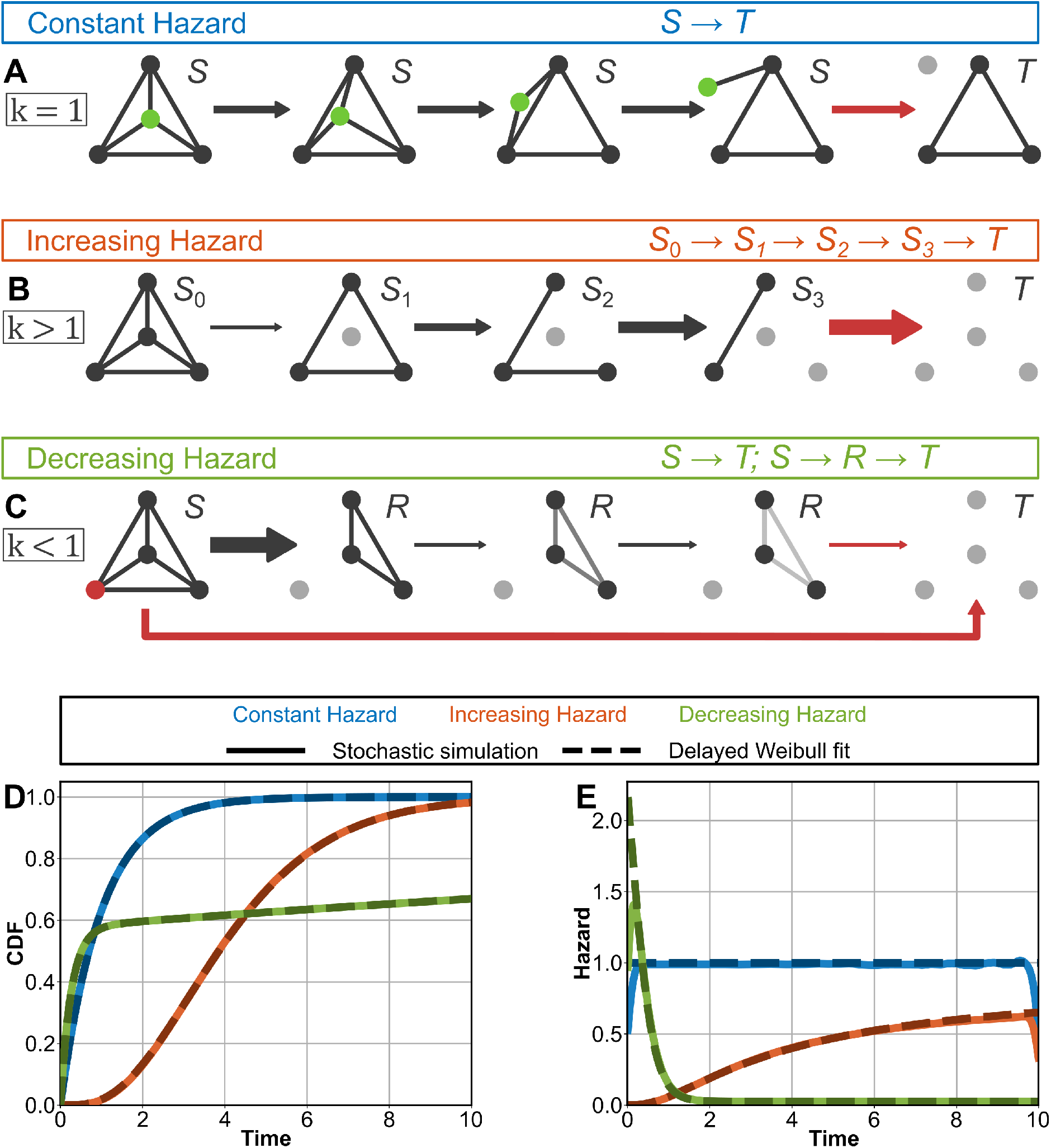
Distinct modes of the shape parameter *k* can emerge from simple classes of network topology. **A-C**. Representative network motifs associated with constant hazard and memoryless escape (**A**), increasing hazard through cumulative destabilisation (**B**), and decreasing hazard through early depletion of the susceptible state (**C**). Each topology was simulated stochastically to generate first-passage-time distributions for cell-state transition. **D**,**E**. Empirical cumulative distribution functions (**D**) and hazard functions (**E**) obtained from these stochastic simulations. Solid lines show stochastic simulation results and dashed lines show delayed Weibull fits (*N* = 10^8^ cells). Together, these examples show that distinct delayed Weibull regimes, and therefore distinct fitted values of *k*, can arise from qualitatively different underlying transition architectures.

When *k >* 1, the hazard increases with time, since

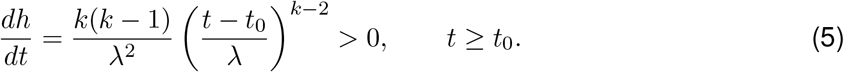

Thus, the longer a cell remains in the pre-transition state after *t*_0_, the greater its instantaneous probability of transition. A minimal topology consistent with this regime is the sequential process

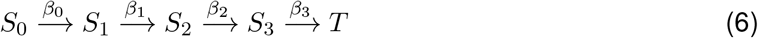

(Fig. 2 B), with *β*_*i*_ *< β*_*i*+1_. For the rates used here, (*β*_0_, *β*_1_, *β*_2_, *β*_3_) = (0.7, 0.9, 1.1, 1.3), stochastic simulation of this topology recovers a delayed Weibull fit with *λ* = 4.667264 and *k* = 2.114808. The resulting cumulative distribution function displays the delayed, progressively steepening profile expected from a multi-step escape process (Fig. 2 D, orange lines). This, in turn, gives rise to a hazard that increases through time (Fig. 2 E, orange lines). This regime is consistent with cumulative or cooperative destabilisation and may arise from positive feedback, progressive loss of redundancy, or a requirement for multiple regulatory constraints to be removed before transition becomes likely.

When *k <* 1, the hazard decreases with time, since

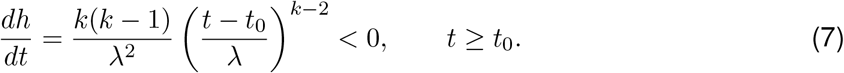

Thus, cells are most likely to transition soon after *t*_0_, with instantaneous transition probability decreasing thereafter. A minimal topology consistent with this regime is the competing-pathway process

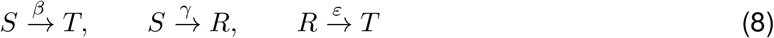

(Fig. 2 C). For the rates used here, *β* = 2.2, *γ* = 1.6, and *ε* = 0.025, stochastic simulation of this topology recovers a delayed Weibull fit with *λ* = 5.035541 and *k* = 0.195215. The resulting cumulative distribution function shows rapid early accumulation followed by pronounced deceleration, consistent with the early depletion of the susceptible state (Fig. 2 D, green lines). This, in turn, gives rise to a hazard that decreases over time (Fig. 2 E, green lines). In this regime, cells that do not transition rapidly from the susceptible state *S* are diverted into a more refractory state *R*, from which subsequent transitions remain possible but occur slowly. Biologically, such behaviour may reflect robust negative feedback, homeostatic buffering, or rapid adaptation away from an initially fragile configuration.

Taken together, these examples demonstrate that distinct modes of *k* can emerge from qualitatively different topologies and transition mechanisms. The shape parameter *k* can therefore be interpreted as a coarse-grained readout of the temporal organisation of state exit. Values near *k* = 1 are consistent with memoryless escape, values *k >* 1 with cumulative or cooperative destabilisation, and values *k <* 1 with early susceptibility followed by refractory stabilisation or depletion of the responsive state. While these minimal models are illustrative rather than exhaustive, they show that fitted values of *k* can provide mechanistically informative clues about the underlying biology of a transition.

### A practical workflow for fitting and comparing delayed Weibull models across experimental designs

Having derived the model and interpreted its parameters, we next considered how it can be fit and compared across experimental conditions. Applying the delayed Weibull model to data requires a sequence of linked modelling decisions: which parameters to estimate or fix, whether to fit replicate time courses separately or jointly, and which statistical comparisons are appropriate for the resulting fits (Fig. 3). The model describes a transition time course using four parameters, the onset delay *t*_0_, the characteristic timescale *λ*, the shape parameter *k*, and the competent fraction *π*. These parameters may be estimated from observed data using a general nonlinear optimization method such as nonlinear least squares. In practice, this can be implemented using standard routines such as `lsqcurvefit` (MATLAB), `least_squares` from `scipy.optimize` (Python), or `nls` (R). The input is typically a time series of observed transitioned fractions (or a suitably normalized equivalent). The output is a fitted parameter vector *θ* = (*t*_0_, *λ, k, π*), from which fitted time courses and derived timing metrics can be obtained, together with approximate uncertainty measures when supported by the fitting procedure.

**Figure 3.**
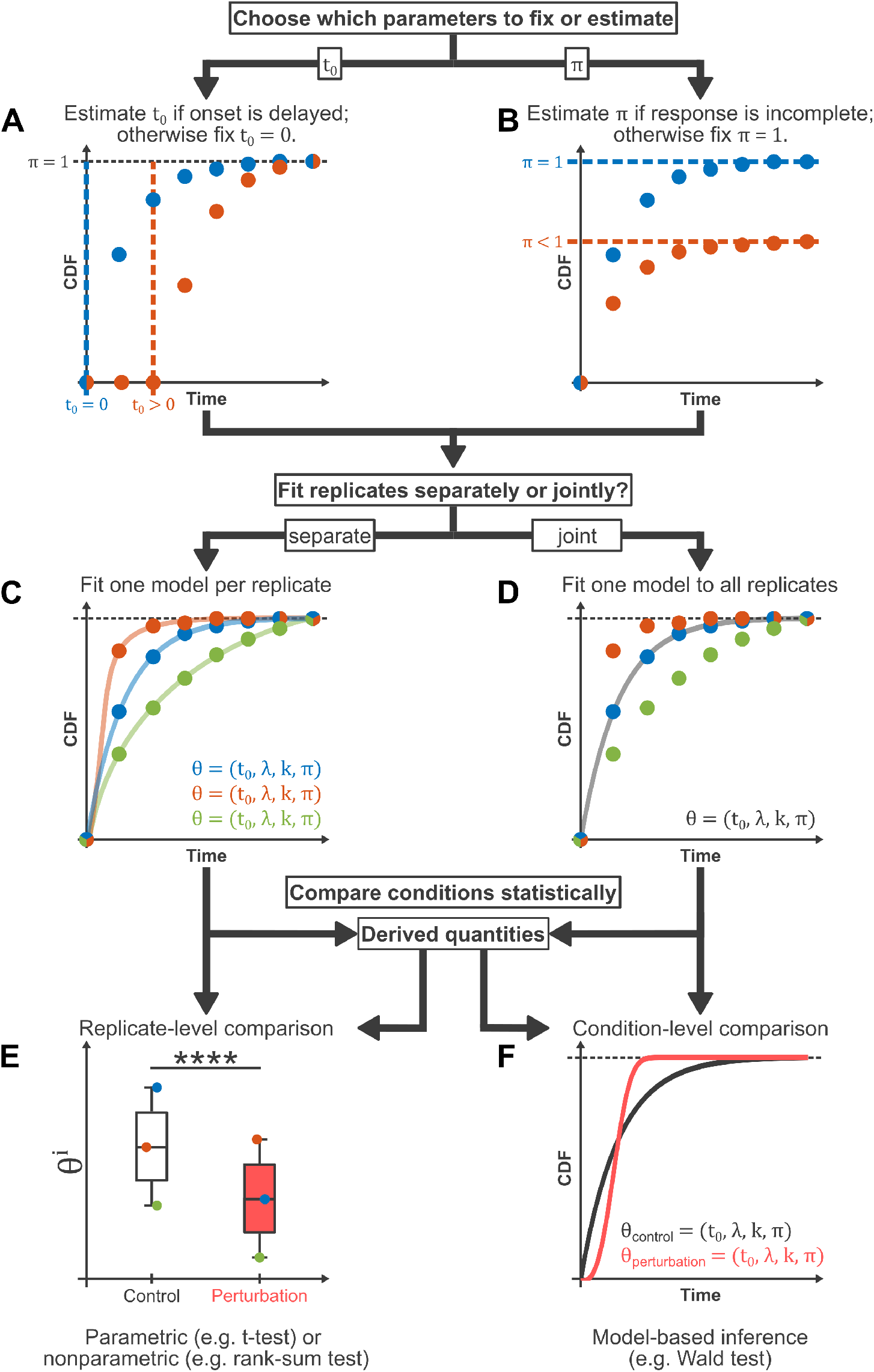
A practical workflow links parameterization, fitting strategy, and statistical inference for delayed Weibull models across experimental designs. **A**,**B**. Selection of which parameters should be fixed or estimated. The onset-delay parameter *t*_0_ may be estimated when transition begins after a clear lag, and otherwise may be fixed to 0 (**A**). The competent fraction *π* may be estimated when the response is incomplete, and otherwise may be fixed to 1 (**B**). **C**,**D**. Alternative fitting strategies depending on the structure of the data. Replicate-resolved time courses may be fit separately with one model per replicate (**C**), whereas aggregated or condition-level time courses may be fit jointly with one model per condition (**D**). **E**,**F**. Corresponding approaches to statistical comparison. Inference may be performed on fitted parameters directly or on quantities derived from them. Separate replicate-level fits permit comparison across replicates using conventional parametric or nonparametric tests (**E**), whereas joint condition-level fits permit model-based inference on fitted parameters or derived quantities such as *t*_50_ (**F**). Together, these choices determine how delayed Weibull models should be fit, interpreted, and compared across conditions.

The onset delay *t*_0_ should be estimated when the data suggest that a transition begins only after a lag - indicated by the measured fraction remaining near zero for an initial period (Fig. 3 A). Conversely, if the transition begins immediately or the sampling interval is too coarse to resolve a lag, fixing *t*_0_ = 0 is often preferable. Similarly, the competent fraction *π* should be estimated if the response is incomplete and saturates below one (Fig. 3 B). If the data are consistent with eventual complete transition, fixing *π* = 1 avoids an unnecessary degree of freedom.

When replicate-resolved time courses are available, fitting one model per replicate is generally the preferred approach for reproducibility. This allows for the explicit quantification of replicate-to-replicate variability (Fig. 3 C) and supports downstream statistical comparison at the replicate level. Jointly fitting a single model to all observations within a condition (Fig. 3 D) may be necessary when data are only available in an aggregated form or material is intrinsically limited or difficult to acquire (e.g. rare primary tissue samples).

The chosen fitting strategy directly determines the appropriate methodology for downstream statistical inference. When replicates are processed separately, comparisons are typically performed on replicate-level parameter estimates or summary statistics, such as the time to 50% transition, *t*_50_, using standard parametric or non-parametric tests (Fig. 3 E). Conversely, when conditions are fit jointly, inference must be performed directly on the fitted parameters and their derived quantities using nonlinear regression inference methods. These include techniques based on local approximation, likelihood regions, and profile-based summaries (Fig. 3 F) (Bates and Watts, 1988).

In these joint-fitting scenarios, differences between conditions are assessed using approximate Wald-type tests (Wald, 1943), likelihood-ratio tests (Wilks, 1938), or extra-sum-of-squares tests for nested model comparisons (Bates and Watts, 1988). To accurately quantify uncertainty, researchers should employ either profile-likelihood methods (Venzon and Moolgavkar, 1988) or bootstrap-based approaches (Efron, 1979, 1987).

From each model, characteristic times, such as *t*_10_, *t*_50_, and *t*_90_, can be calculated analytically. These provide interpretable summaries of timing differences that complement raw parameters. Goodness of fit may be summarised using quantities such as *R*^2^; however, when model comparison is the primary goal, information criteria such as AIC or BIC are preferable (Akaike, 1974; Schwarz, 1978).

This workflow provides a flexible framework for the delayed Weibull model across diverse experimental designs. To demonstrate its utility, we applied it to three systems differing in both biology and measurement modality: bulk gene-expression loss during exit from pluripotency; single-cell lineage acquisition during directed differentiation; and clonal well-based readouts of cellular reprogramming. These examples illustrate the generality of the framework and the specific constraints that different measurements place on biological interpretation.

### Bulk qPCR reveals distinct modes of pluripotency-factor downregulation during exit from self-renewal

In Leeb et al. 2014, haploid mouse embryonic stem cells were maintained in self-renewal conditions (2i) and then transferred to N2B27 to induce exit from the naïve state (Fig. 4 A). Transcript levels of *Esrrb, Klf2, Klf4, Nanog, Tbx3*, and *Tfcp2l1* were measured by RT-qPCR over a 48 h time course. The study also examined the effect of *Pum1* knockdown on the same transition, reporting that *Pum1* depletion delayed downregulation of multiple naïve pluripotency-associated transcripts. In our reanalysis, the time-course data from Figure 3 F and Supplementary Figure S3 C of Leeb et al. 2014 were digitized and fitted with the delayed Weibull model.

**Figure 4.**
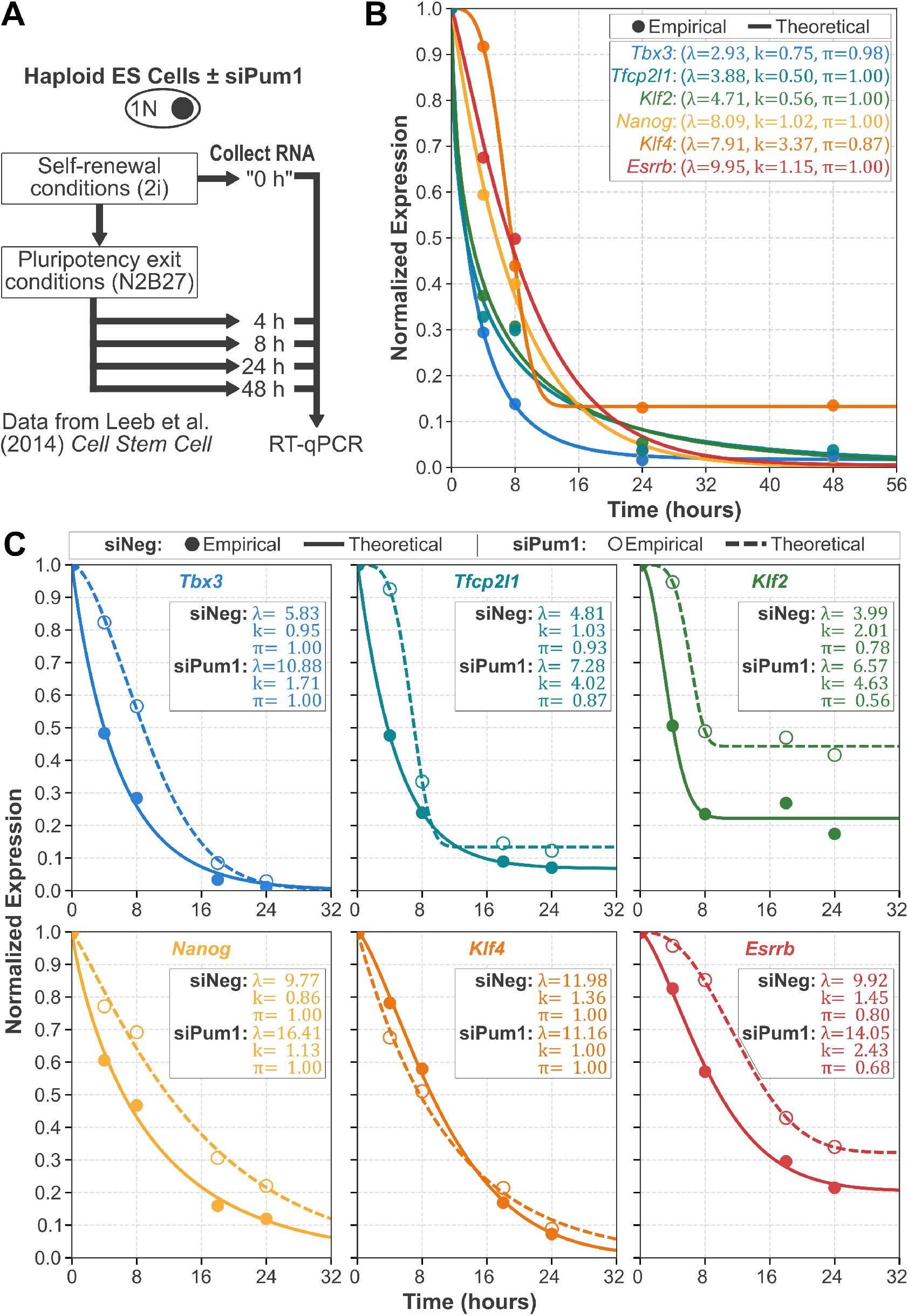
The delayed Weibull model resolves distinct modes of bulk pluripotency-factor down-regulation during exit from naïve pluripotency. **A**. Experimental scheme for Leeb et al. (2014). Haploid mouse embryonic stem cells were maintained in self-renewal conditions (2i), transferred to N2B27 to induce exit from naïve pluripotency, and sampled over a 48 h time course for RT-qPCR. For the perturbation analysis, cells were transfected with control siRNA or si*Pum1* prior to induction of exit. Data represent relative gene expression measured by RT-qPCR and normalized to the expression level at 0 hours. **B**. Bulk down-regulation kinetics of key naïve pluripotency transcription factors during exit from the naïve state resolve an early group (*Tbx3, Tfcp2l1*, and *Klf2*) and a later group (*Nanog, Klf4*, and *Esrrb*), with *Klf4* showing especially sharp down-regulation. Filled circles show empirical data from Figure 3 F of Leeb et al. (2014), and solid lines show delayed Weibull model fits performed here. **C**. *Pum1* knockdown slows or limits down-regulation of multiple naïve pluripotency-associated transcripts during exit from naïve pluripotency. Filled circles and solid lines denote siNeg, whereas open circles and dashed lines denote si*Pum1*. Data are from Supplementary Figure S3 C of Leeb et al. (2014), and fits were performed here.

A limitation of these data is their acquisition from bulk RT-qPCR rather than at single-cell resolution. Consequently, they report only the population-averaged loss of marker expression and cannot distinguish between asynchronous all-or-none switching, gradual cohort-wide downregulation, or a combination of these behaviors. We therefore interpret these trajectories as bulk transition profiles. For these fits, the onset delay was fixed at *t*_0_ = 0, while *λ, k*, and *π* were estimated; this allowed the asymptotic level of downregulation for each marker to be determined from the data rather than assuming it approached zero.

The delayed Weibull model accurately captured the bulk downregulation kinetics for all six genes (Fig. 4 B), with all fits achieving *R*^2^ > 0.98. Two temporal groups were apparent: *Tbx3, Tfcp2l1*, and *Klf2* were downregulated earliest, with fitted *t*_50_ values of 1.80, 1.87, and 2.46 h, respectively; whereas *Nanog, Klf4*, and *Esrrb* responded later, with *t*_50_ values of 5.65, 7.09, and 7.24 h. This divides the naïve pluripotency network into an early-responding group and a late-responding group. Most genes showed relatively shallow or near-exponential declines, with *k* ranging from 0.50 to 1.15; however, *Klf4* was distinct, exhibiting a substantially higher synchronicity parameter (*k* = 3.37). Thus, although *Nanog* and *Klf4* had similar characteristic timescales (*λ* = 8.09 and 7.91, respectively), the more synchronous decline of *Klf4* shifted its *t*_50_ later. Detailed parameter estimates and pairwise statistical comparisons are provided in Supplemental Table S1.

For the *Pum1* perturbation, the fitted trajectories in Figure 4 C recapitulated the qualitative conclusion of the original study that *Pum1* promotes timely dismantling of the naïve pluripotency program. Knockdown of *Pum1* increased the fitted characteristic timescale *λ* for *Tbx3, Tfcp2l1, Klf2, Nanog*, and *Esrrb*, indicating slower transcript loss across much of the network, whereas *Klf4* changed little. In several cases, particularly *Klf2* and *Esrrb, Pum1* knockdown also reduced the fitted value of *π*, consistent with less complete downregulation within the observation window. Together, these fits show that the Weibull framework usefully separates three features of bulk gene-expression decline during exit from self-renewal: overall pace (*λ*), temporal sharpness (*k*), and extent of response (*π*).

### Single-cell measurements reveal how starting state shapes directed differentiation kinetics

In the study by Mulas et al. (2017), the authors examined how mouse embryonic stem cells acquire competence for somatic lineage specification as they exit the naïve state. Differentiation was initiated from three defined starting populations (Fig. 5 A). The first consisted of cells maintained in conventional two-inhibitor medium (2i), which preserves naïve pluripotency (Ying et al., 2008). The other two were obtained by releasing cells from 2i into N2B27 - the basal medium used in 2i culture, which does not support the pluripotent state - for 24 h and then sorting them into Rex1-High and Rex1-Low fractions based on the expression of the naïve pluripotency-associated reporter Rex1::GFPd2. Rex1 is a marker of undifferentiated embryonic stem cells that is downregulated during exit from naïve pluripotency (Toyooka et al., 2008; Kalkan et al., 2017). These three populations were then subjected to directed differentiation towards primitive streak, neural, lateral mesoderm, and definitive endoderm fates. In these assays, the measured readout is the acquisition of the new state rather than the direct loss of the original one. Accordingly, applying the delayed Weibull framework here assumes that the loss of the preceding identity is sufficiently coupled to the acquisition of the new identity that the two can be treated as a single effective transition at the phenotypic level. In our reanalysis, the time courses for primitive streak, neural, lateral mesoderm, and definitive endoderm differentiation were digitized from Figure 2A, K, C, and E of Mulas et al. (2017), respectively, and fitted with the delayed Weibull model.

**Figure 5.**
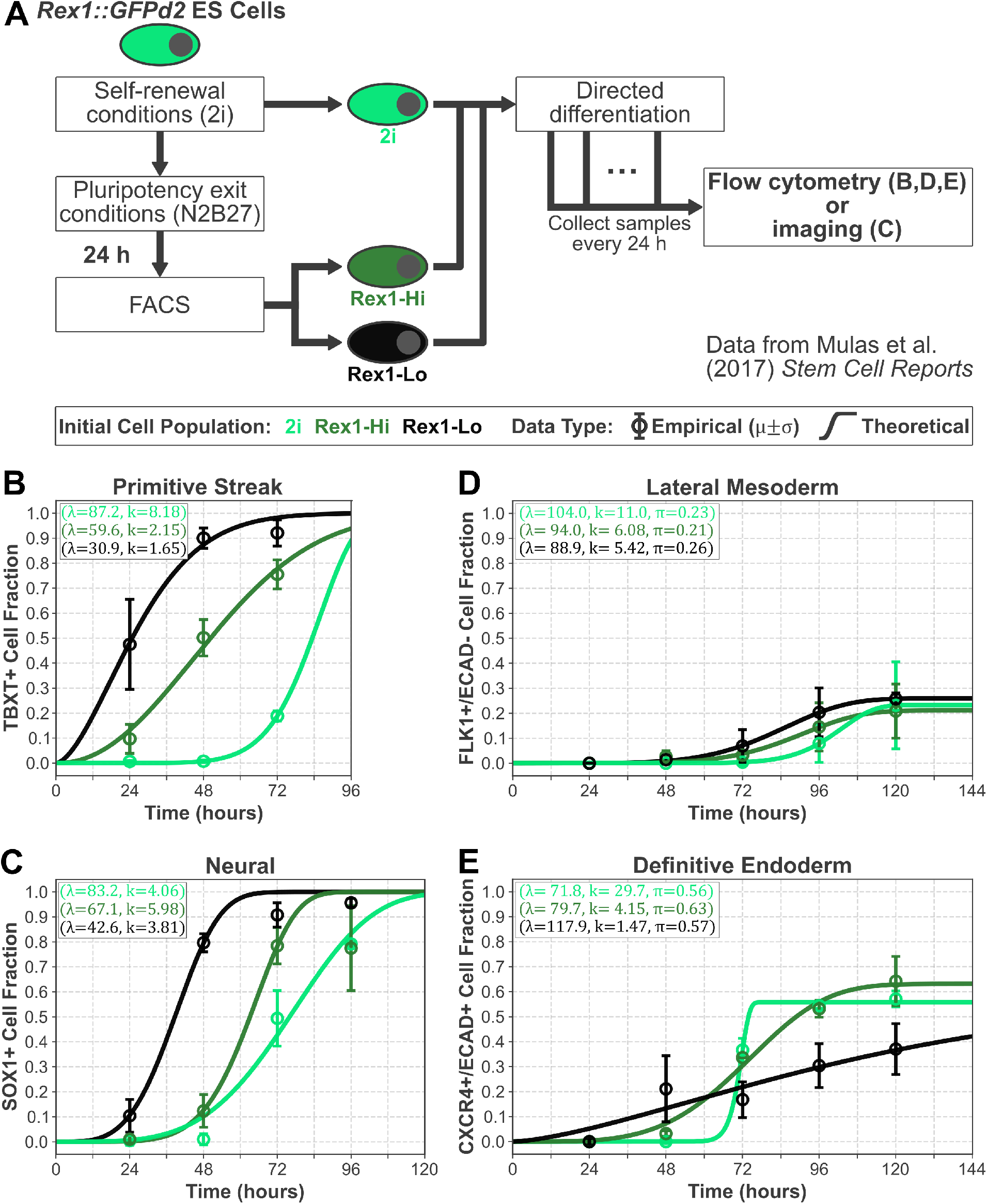
The delayed Weibull model quantifies how starting state shapes the timing, synchronicity, and extent of directed differentiation. **A**. Experimental scheme for Mulas et al. (2017). Cells were maintained in 2i or released into N2B27 for 24 h and then sorted on the basis of Rex1::GFPd2 expression before directed differentiation. Differentiation was therefore initiated from three starting populations: 2i (light green), Rex1-Hi (dark green), and Rex1-Lo (black). Primitive streak, lateral mesoderm, and definitive endoderm were assayed by flow cytometry, whereas neural differentiation was assayed by immunostaining and imaging. **B–E**. Directed differentiation toward primitive streak (**B**; data from Figure 2 A of Mulas et al. (2017)), neural lineage (**C**; data from Figure 2 K of Mulas et al. (2017)), lateral mesoderm (**D**; data from Figure 2 C of Mulas et al. (2017)), and definitive endoderm (**E**; data from Figure 2 E of Mulas et al. (2017)). Open circles with error bars show empirical mean *±* standard deviation, and solid lines show delayed Weibull model fits performed here. Primitive streak and neural differentiation show the clearest progression toward earlier response in Rex1-Lo cells relative to Rex1-Hi and 2i cells. Lateral mesoderm and definitive endoderm both show incomplete differentiation competence, but only lateral mesoderm reveals clear differences in responding fraction between starting states.

We applied the delayed Weibull model to these data to quantify and compare differentiation dynamics across the three starting populations. For primitive streak and neural differentiation, we fitted *λ* and *k*, with *π* fixed to 1, because nearly the entire population acquired marker expression. For lateral mesoderm and definitive endoderm, we additionally fitted *π* to capture the reduced competent fraction apparent in these assays. Across all lineages, the model captured the observed trajectories well, with all but one fit having *R*^2^ > 0.9 (Fig. 5 B–E). Under the assumption that the loss of the preceding state is tightly coupled to the installation of the new one, earlier acquisition of lineage-marker expression is interpreted as earlier effective completion of the transition. A consistent trend was that cells further from the naïve state generally completed this effective transition earlier than cells maintained in 2i. This was most evident in the primitive streak and neural assays (Fig. 5 B,C), where Rex1-Lo cells acquired lineage-marker expression earliest, Rex1-Hi cells were intermediate, and 2i cells were slowest.

For primitive streak differentiation (Fig. 5 B), the fitted characteristic timescale decreased progressively from 2i (*λ* = 87.2) to Rex1-Hi (*λ* = 59.6) to Rex1-Lo (*λ* = 30.9). The corresponding *t*_50_ values showed the same ordering, consistent with the close relationship between *λ* and the more readily interpretable *t*_50_. The 2i population also showed the most synchronous transition, with a substantially larger fitted *k* value (*k* = 8.18) than either Rex1-Hi (*k* = 2.15) or Rex1-Lo (*k* = 1.65). However, this apparent increase in synchronicity should be interpreted cautiously, as the relatively sparse sampling of the trajectory, particularly for the slower 2i condition, may inflate the fitted value of *k*. Thus, within the sampled time window, progression out of naïve pluripotency appeared to accelerate primitive streak induction, but stronger inference about differences in temporal coordination would require a longer time course that more fully resolves the 2i response.

Neural differentiation showed a similar ordering in timing (Fig. 5 C). The fitted *λ* values again decreased from 2i (*λ* = 83.2) to Rex1-Hi (*λ* = 67.1) to Rex1-Lo (*λ* = 42.6), and Rex1-Lo cells reached *t*_50_ earlier than either of the other two populations. In contrast to primitive streak differentiation, however, the fitted *k* values were similar across all three starting states. This suggests that neural induction becomes faster as cells leave the naïve state, while the sharpness of the transition remains broadly comparable.

For lateral mesoderm differentiation (Fig. 5 D), the same temporal gradient was retained, but with a lower overall responding fraction. The fitted timescale again decreased from 2i (*λ* = 104.0) to Rex1-Hi (*λ* = 94.0) to Rex1-Lo (*λ* = 88.9). Both Rex1-Hi and Rex1-Lo therefore differentiated earlier than 2i cells, while remaining similar to each other. The 2i population again displayed the most synchronous dynamics (*k* = 11.0) relative to Rex1-Hi (*k* = 6.08) and Rex1-Lo (*k* = 5.42). In this assay, the fitted competent fraction was highest for Rex1-Lo (*π* = 0.26), intermediate for 2i (*π* = 0.23), and lowest for Rex1-Hi (*π* = 0.21). Thus, release from the naïve state was associated not only with faster lateral mesoderm induction but also with a modest increase in the responding fraction.

Definitive endoderm differed from the other lineages (Fig. 5 E). Although the fits remained strong for 2i and Rex1-Hi, and slightly weaker for Rex1-Lo (*R*^2^ = 0.8584), this case highlights an important limitation of the framework. Mulas et al. (2017) reported high levels of cell death in the Rex1-Lo population under these induction conditions. In such cases, the observed trajectory is confounded, because the measured signal may reflect not only the kinetics of state transition but also differential survival of the responding population. Consequently, fitted parameters may capture a mixture of transition dynamics and cell-loss effects, rather than transition kinetics alone. This example illustrates that conditions confounded by significant cell death - or by other processes that substantially distort the population trajectory - must be controlled for or else treated as limits of interpretation for the model. For this reason, the absence of clear differences in fitted *t*_50_, *k*, or *π* in this instance should not be over-interpreted.

Taken together, these analyses support a general picture in which exit from the naïve state increases readiness to respond to lineage-inductive cues. Rex1-Lo cells were typically the fastest to upregulate lineage markers, Rex1-Hi cells were intermediate, and 2i cells were slowest. At the same time, the 2i population often showed equal or greater synchronicity of differentiation, consistent with a more homogeneous starting state. These findings suggest that progression from naïve pluripotency promotes lineage responsiveness while also increasing heterogeneity in the timing of transition across the population. Full parameter estimates and pairwise statistical comparisons are provided in Supplemental Table S2.

### Clonal well-based assays distinguish perturbations that accelerate reprogramming after population-level rescaling

In Hanna et al. 2009, monoclonal pre-B-cell populations carrying a secondary doxycycline-inducible OSKM reprogramming system and a Nanog-GFP reporter were followed over time to quantify the acquisition of pluripotency (Fig. 6 A). The parental NGFP1 line was compared with derivative lines carrying additional perturbations: Lin28 overexpression, Nanog overexpression, p21 knockdown, or p53 knockdown. Single transgenic pre-B cells were sorted into individual wells, cultured in doxycycline, and screened weekly for reactivation of the endogenous Nanog-GFP reporter. As Nanog is a key transcription factor associated with the naïve pluripotent state, reactivation of the Nanog-GFP reporter served as a readout for successful reprogramming. A threshold of > 0.5% Nanog-GFP^+^ cells per well was defined as successful reprogramming, permitting the derivation of stable doxycycline-independent iPSC lines. In this assay, the measured readout is the acquisition of a new state rather than the direct loss of the original one. Accordingly, applying the delayed Weibull framework here assumes that the collapse of the starting state is sufficiently coupled to the establishment of the pluripotent state to be treated as a single effective transition. For our reanalysis, the population-rescaled time courses for the parental NGFP1 line and its derivatives were digitized from Figure 4d of Hanna et al. (2009) and fitted with the delayed Weibull model.

**Figure 6.**
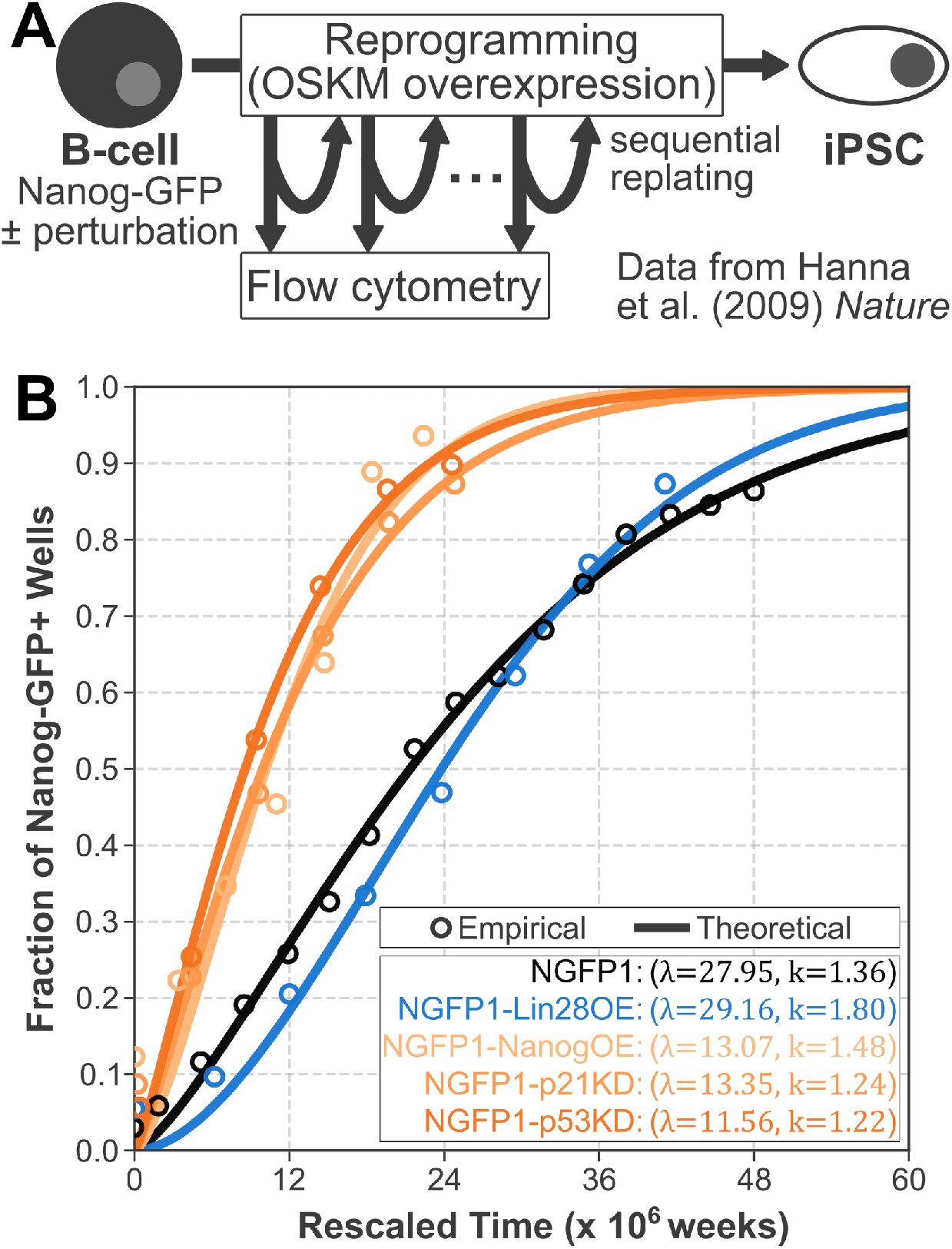
The delayed Weibull model separates perturbations that accelerate reprogramming after population-level time rescaling. **A**. Experimental scheme for Hanna et al. (2009). Secondary transgenic pre-B cells carrying doxycycline-inducible OSKM factors and a Nanog-GFP reporter were plated as clonal populations, cultured in doxycycline, and screened sequentially for the appearance of Nanog-GFP^+^ wells during reprogramming. **B**. Reprogramming kinetics of monoclonal pre-B-cell populations for the parental NGFP1 line and derivative lines carrying Lin28 over-expression (OE), Nanog OE, p21 knockdown (KD), or p53 KD. Open circles show empirical data from Figure 4 d of Hanna et al. (2009), and solid lines show delayed Weibull model fits performed here. After population-level time rescaling, Nanog OE, p21 KD, and p53 KD remain clearly left-shifted relative to NGFP1, whereas Lin28 OE remains close to the parental trajectory.

The original study interpreted these data using a stochastic framework where the cumulative appearance of Nanog-GFP^+^ wells was analyzed after rescaling time by effective population size and proliferation rate. This representation distinguished perturbations that primarily accelerated reprogramming through increased proliferation from those that altered the underlying cell-intrinsic rate. Specifically, the original analysis concluded that p53/p21 inhibition and Lin28 overexpression acted predominantly through cell-division-rate-dependent effects, whereas Nanog overexpression accelerated reprogramming in a largely cell-division-rate-independent manner.

We reanalyzed these population-rescaled time courses using the delayed Weibull model, with *t*_0_ fixed to 0 and *π* fixed to 1, such that *λ* and *k* captured the characteristic timescale and temporal sharpness of reprogramming, respectively (Fig. 6 B). Under the assumption that B-cell state loss is tightly coupled to the acquisition of pluripotency, earlier Nanog-GFP activation reflects an earlier effective completion of reprogramming. The model provided a robust description of the cumulative Nanog-GFP activation curves across all conditions (*R*^2^ > 0.92; Fig. 6 B). Three perturbations - Nanog overexpression, p21 knockdown, and p53 knockdown - were associated with substantially shorter characteristic timescales than the parental NGFP1 line (*λ* = 13.07, 13.35, and 11.56, respectively, vs *λ* = 27.95 for NGFP1). The corresponding fitted times to 50% reprogramming (*t*_50_) were 10.20, 9.92, and 8.57, compared with 21.35 for NGFP1 (in units of population-rescaled time). By contrast, NGFP1-Lin28OE remained close to the parental condition after rescaling, with *λ* = 29.16 and *t*_50_ = 23.79. Thus, in this representation, Nanog overexpression and p53/p21 inhibition remained clearly left-shifted relative to NGFP1, whereas Lin28 overexpression did not.

Differences in the synchronicity parameter (*k*) were comparatively modest. NGFP1-Lin28OE showed the highest fitted value (*k* = 1.80), indicating a slightly sharper transition than the other conditions, which ranged from *k* = 1.22 (NGFP1-p53KD) to *k* = 1.48 (NGFP1-NanogOE). Overall, this reanalysis separates the perturbations into two phenotypic groups in population-rescaled time: Nanog overexpression and p53/p21 knockdown accelerated the appearance of Nanog-GFP^+^ wells, whereas Lin28 overexpression followed the parental NGFP1 trajectory once proliferation-dependent effects were normalized. Detailed parameter estimates and pairwise statistical comparisons are provided in Supplemental Table S3. Together, these results show that, after accounting for population-level proliferation effects, the delayed Weibull framework distinguishes perturbations that genuinely accelerate effective reprogramming from those with weaker residual effects on transition timing.

## DISCUSSION

We have presented a compact parametric framework for quantifying cell-state transition kinetics by treating state exit as the cumulative consequence of many stochastic molecular events - analogous to failure in a stabilising regulatory network. This yields a delayed Weibull description with four interpretable parameters: an onset delay *t*_0_, a characteristic timescale *λ*, a shape parameter *k* describing the temporal hazard, and a competent fraction *π*. Together, these parameters decouple distinct features of a transition that are often conflated in qualitative descriptions of time-course data.

This framework provides a common language for comparing transitions across diverse biological settings. We demonstrated that the same formalism describes the loss of pluripotency-associated transcripts during exit from self-renewal, the gain of lineage markers during directed differentiation, and the acquisition of a pluripotency reporter during reprogramming. These systems vary not only in their underlying biology but also in their measurement modalities: bulk RT-qPCR reports population averages; flow cytometry and imaging report marker-positive fractions; and plate-based assays report the cumulative fraction of wells undergoing a detectable event. That the delayed Weibull form captures these disparate datasets suggests that it reflects a generic statistical structure of cell-state transitions, regardless of the molecular details.

However, the interpretation of the fitted parameters must be conditioned on experimental design. For bulk measurements, the model describes the kinetics of change in the population-average signal but cannot distinguish between discrete switching and gradual change. For flow cytometry and imaging, the curves more directly represent transition fractions, whereas for plate-based assays, the observable is a clonal detection event rather than an individual cell state. More generally, when the assay measures acquisition of a new state rather than loss of the original one, the present framework should be interpreted as describing an effective transition whose timing reflects the degree of coupling between network collapse and network installation.

Among the parameters, *k* is especially informative, as it moves beyond a simple summary of speed to report how transition propensity changes over time. Values near *k* = 1 correspond to an approximately memoryless process, *k >* 1 to increasing transition propensity, and *k <* 1 to early susceptibility followed by decreasing hazard. The minimal topological examples in Figure 2 show that these regimes can emerge from qualitatively distinct classes of network organisation. Nevertheless, we caution against over-interpretation: sparse sampling, limited dynamic range, and imperfect observability can all influence *k*. Thus, it should be viewed as a potentially informative coarse-grained descriptor rather than a direct mechanistic readout.

The competence parameter *π* is similarly valuable, separating the extent of a response from its kinetics. This is particularly useful when transitions remain incomplete within the observation window; here, the model distinguishes a reduced responding fraction from a slower transition among competent cells. These outcomes are biologically distinct but rarely separated explicitly in descriptive analyses.

The framework is deliberately lightweight. Unlike high-dimensional trajectory models or explicit regulatory-network reconstructions, the delayed Weibull model can be fitted to relatively small time-course datasets using standard nonlinear least-squares methods. It is therefore well-suited to legacy data, low-dimensional assays, and systems where dense temporal sampling is impractical. Its purpose is to complement, rather than replace, mechanistic or single-cell approaches by providing a compact quantitative summary of transition kinetics.

This simplicity is a strength, because it makes the framework broadly applicable to datasets in which more detailed modeling would be difficult or unwarranted. At the same time, it defines the present scope of interpretation. The model is descriptive rather than mechanistic, so a good fit alone does not identify which molecular components change, in what order they do so, or whether similar coarse-grained kinetics could arise from different underlying network architectures. It also assumes a single dominant transition process and may therefore average over genuinely distinct kinetic subpopulations. In addition, because *π* is defined relative to the observation window and chosen marker, non-responding cells may include cells that transition later or via alternative routes not captured by the assay. However, when applied in conjunction with careful experimental design and informative sets of perturbations, the framework may help reveal these underlying features indirectly by showing how timing, synchronicity, and response extent shift across conditions.

These considerations also suggest several useful directions for extending and refining the framework beyond the scope of the present study. These include testing its performance on replicate-resolved single-cell datasets and in systems with denser temporal sampling, comparing the delayed Weibull systematically with alternative parametric families, and relating fitted parameters to explicit models of regulatory-network dynamics. Such efforts would help define more clearly where the present formulation is most informative and where more detailed modelling is warranted. Critically, this work argues that the kinetics of cell-state change should be treated as a quantitative phenotype in its own right. By providing a practical means to measure timing, synchronicity, and response extent, the delayed Weibull model offers a robust way to compare transitions across empirical and theoretical experiments and to guide deeper investigation.

## METHODS

### Data collection

Data were collated from published figures and quantified using GetData Graph Digitizer (http://getdata-graph-digitizer.com). Axes were calibrated manually for each panel, and point capture mode was used to extract mean values and, where available, corresponding measures of variation. For datasets in which error bars were shown, standard deviations were digitized when possible. Digitized values were then reformatted into analysis-ready tables for model fitting.

### Model fitting and statistical analysis

All model fitting and statistical analyses were performed in Python. Time courses were fit with the delayed Weibull model using nonlinear least-squares optimization, with parameters fixed or estimated according to the structure of each dataset, as described in the main text. Derived timing metrics, including *t*_10_, *t*_50_, and *t*_90_, were calculated analytically from the fitted parameters. Where condition-level fits were compared directly, pairwise comparisons were performed using approximate Wald-type tests on fitted parameters or derived quantities, with Benjamini–Hochberg correction applied to account for multiple testing. Goodness of fit was summarized using *R*^2^.

## Data and code availability

Digitized datasets and code used for simulations, model fitting, and statistical analysis are publicly available on GitHub (https://github.com/stanleystrawbridge/strawbridge_2026_parametric).

## ACKNOWLEDGMENTS

S.E.S. was supported by a Sir Henry Wellcome Postdoctoral Fellowship (224070/Z/21/Z) and a University of Sheffield Strategic Research Fellowship in the Physics of Life and Quantitative Biology. The authors thank Austin Smith for helpful feedback on the manuscript.

## AUTHOR CONTRIBUTIONS

Conceptualization, S.E.S.; Methodology, S.E.S. and A.G.F.; Software, S.E.S.; Validation, S.E.S. and A.G.F.; Formal analysis, S.E.S. ; Investigation, S.E.S.; Data Curation, S.E.S.; Writing - Original Draft, S.E.S. and A.G.F; Writing - Review & Editing, S.E.S. and A.G.F; Visualization, S.E.S.; Funding acquisition, S.E.S.

## SUPPLEMENTAL MATERIAL

### 1 SUPPLEMENTAL TABLES

#### Supplemental Table S1

Supplemental Table S1 is provided as an XLSX file containing two sheets associated with Fig. 4. The sheet titled “`leeb_2014_cell-stem-cell`” contains the data extracted from Leeb et al. (2014). Figure 3F is reproduced here as Fig. 4 B and Fig. S3 C is reproduced here as Fig. 4 C. The sheet titled “`fitting_&_statistics`” contains the fitting summary and pairwise statistical comparisons for the downregulation of gene expression during exit from naïve pluripotency in Fig. 4 B and the effect of *Pum1* inhibition by RNA interference on the expression patterns of the same genes during exit from naïve pluripotency in Fig. 4 C.

#### Supplemental Table S2

Supplemental Table S2 is provided as an XLSX file containing two sheets associated with Fig. 5. The sheet titled “`mulas_2017_stem-cell-reports`” contains the data extracted from Mulas et al. (2017). Figure 2 A is reproduced here as Fig. 5 B, Fig. 2 K is reproduced here as Fig. 5 C, Fig. 2 C is reproduced here as Fig. 5 D, and Fig. 2 E is reproduced here as Fig. 5 E. The sheet titled “`fitting_&_statistics`” contains the fitting summary and pairwise statistical comparisons for the kinetics of directed differentiation from different starting populations of mouse embryonic stem cells, as determined by culture condition and Rex1::GFPd2 status.

#### Supplemental Table S3

Supplemental Table S3 is provided as an XLSX file containing two sheets associated with Fig. 6. The sheet titled “`hanna_2009_nature`” contains the data extracted from Hanna et al. (2009). Figure 4 d is reproduced here as Fig. 6 B. The sheet titled “`fitting_&_statistics`” contains the fitting summary and pairwise statistical comparisons for the reprogramming efficiency of B cells to the pluripotent state, with and without genetic perturbations that may affect reprogramming efficiency.

## 2 SUPPLEMENTAL MODEL

### 2.1 A failure-based framework yields a delayed Weibull description of cell-state transition kinetics

We consider a cell state maintained by a stabilising regulatory network composed of *N* regulatory nodes. Each node represents a molecular component, such as a transcription factor, chromatin state, signalling pathway, or mechanical input, that contributes to the stability of the current cellular identity. The state is assumed to be metastable, such that stochastic fluctuations and regulatory turnover may eventually destabilise the network sufficiently to permit transition to an alternative state (Strawbridge et al., 2026).

Inspired by Weibull (1951), for each node *i* ∈ {1, …, *N* } we define the failure time, *τ*_*i*_, as a continuous random variable representing the time at which that node ceases to support the current state (Fig. 1A). Here, ‘failure’ may correspond to degradation, silencing, or functional inactivation. We denote the cumulative distribution function of *τ*_*i*_ by *P*_*i*_(*t*) = ℙ(*τ*_*i*_ ≤ *t*).

For each node, the failure time *τ*_*i*_ is described by a time-dependent hazard *h*_*i*_(*t*), so that

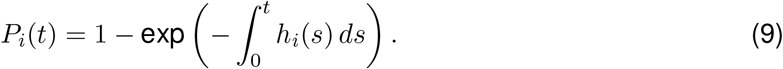

To obtain a tractable expression for the behaviour of the full network, we make the simplifying assumption that node failures are conditionally independent given the state of the system. We also assume that the probability of failure over a short interval is small.

We now consider the probability that the regulatory network has not yet undergone a state transition by time *t*. This survival probability is denoted by *S*(*t*). A transition is assumed to become increasingly likely as destabilisation of the network accumulates. Under the additional assumptions that individual failure contributions combine additively and that the probability of multiple simultaneous failures is negligible, the total cumulative hazard of the network may be written as

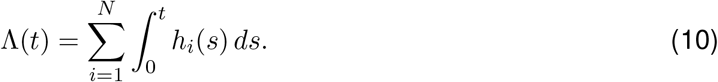

The corresponding survival probability of the network is then

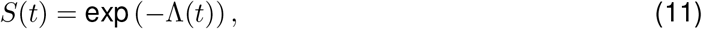

yielding a cumulative distribution function of transition times of

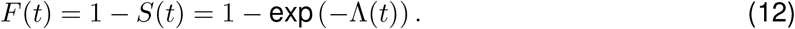

Motivated by our previous analysis of exit from naïve pluripotency, in which a progressively time-dependent failure process provided a useful description of state-transition kinetics (Strawbridge et al., 2026), we now adopt a more general coarse-grained assumption for node failure within the stabilising regulatory network. Specifically, after a common onset delay *t*_0_, we assume that the instantaneous propensity for failure of each node varies with elapsed time. Under cumulative destabilisation, it is natural to expect that this propensity will often increase with time, although constant or decreasing failure propensities are also admissible special cases. As a minimal form that captures these possibilities while remaining analytically tractable, we assume that the hazard for each node follows a shared power-law dependence on elapsed time, differing only by a node-specific scaling factor. Thus, for *t* ≥ *t*_0_, we write

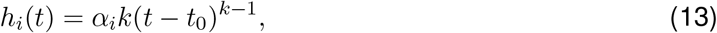

where *α*_*i*_ > 0 is a node-specific scaling parameter, *k >* 0 is a shared shape exponent, and *t*_0_ ≥ 0 is a common onset delay. Integrating this hazard gives

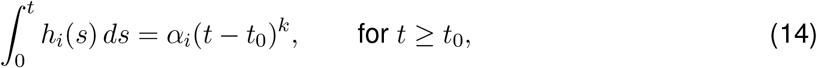

so that the cumulative contribution of each node also follows a shared power-law form. This assumption is appropriate when nodes fail through a common underlying destabilisation process, such that the temporal form of the hazard is shared across nodes while differences in their contributions are captured by the scaling parameters *α*_*i*_. In this case, introducing the scale parameter

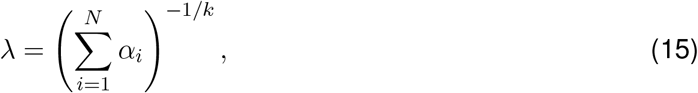

we may write the cumulative distribution function for the competent subpopulation, *F*_c_(*t*), in standard delayed Weibull form:

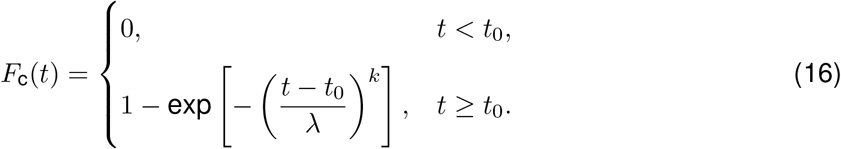

To account for the fact that not all cells in a population are competent to undergo the transition, we introduce a competence parameter *π* ∈ [0, 1]. This parameter denotes the fraction of cells that are capable of responding and eventually transitioning, as judged, for example, by downregulation of one marker or upregulation of another. The remaining fraction is assumed to be non-responding over the timescale of observation.

The cumulative distribution function for the full population is therefore given by

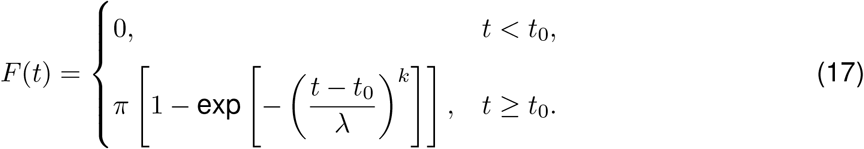

Thus, under the assumptions of stochastic node failure, additive accumulation of destabilisation, a finite onset delay, and partial population competence, the time to cell-state transition is expected to follow a delayed Weibull distribution scaled by the competent fraction *π*.

### 2.2 The delayed Weibull parameters separate onset delay, pace, temporal hazard, and response extent

This formulation separates four distinct features of the population response. The parameter *t*_0_ defines the onset delay, *λ* sets the characteristic timescale of transition among competent cells, *k* determines the temporal form of the transition-time distribution, and *π* specifies the fraction of cells that are able to execute the transition within the observation window.

The delay parameter *t*_0_ acts as a pure temporal shift, moving the distribution to later times without changing its shape (Fig. 1 B). An increase in *t*_0_ postpones the onset of transition uniformly across the population, but does not alter the degree of cell-to-cell variability once transition has become possible. Biologically, *t*_0_ may therefore be interpreted as a latency period, refractory interval, or the minimum time required for competence, signal integration, or preliminary molecular reorganisation before a transition can occur.

The scale parameter *λ* stretches the time axis and sets the overall pace of transition (Fig. 1 C). Larger values of *λ* correspond to slower erosion of the stabilising network and hence slower commitment dynamics, whereas smaller values correspond to more rapid progression toward transition.

The most direct interpretation of the shape parameter *k* is through the hazard function for competent cells. Writing

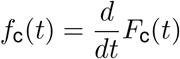

for the corresponding probability density function, the hazard is

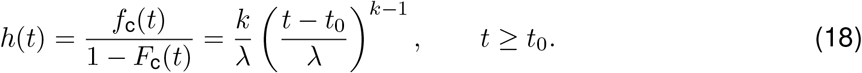

Accordingly, *k* determines whether the instantaneous probability of transition increases, remains constant, or decreases with time after the onset delay (Fig. 1 D). When *k* = 1, the hazard is constant in time, corresponding to a memoryless process with no effective temporal reinforcement (Fig. 1 E). When *k >* 1, the hazard increases with time, so that cells become progressively more likely to transition the longer they remain in the pre-transition state. When *k <* 1, the hazard decreases with time, so that transition is most likely soon after the delay period and less likely thereafter. Potential mechanistic interpretations of these different modes is considered in the section “**Origins of the distinct modes of** *k*”.

The competence parameter *π* controls the asymptotic fraction of the population that undergoes transition,

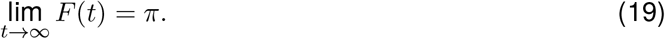

Thus, *π* determines the overall extent of response at the population level rather than the timing of transition among cells that are able to respond (Fig. 1F).

Taken together, these four parameters separate distinct features of the transition process. The parameter *t*_0_ specifies when transitions can begin, *λ* sets the characteristic timescale of transition, *k* determines how transition propensity changes with time after onset, and *π* specifies the fraction of cells that are competent to respond within the observation window.

### 2.3 Moments and characteristic times of the delayed Weibull transition-time distribution

To connect the fitted model to experimentally interpretable timing summaries, we next derived the moments of the delayed Weibull distribution for the competent subpopulation. For the competent subpopulation, the cumulative distribution function of transition times is

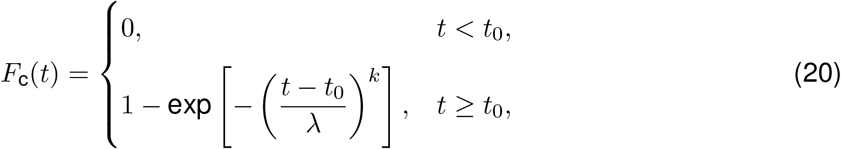

with corresponding probability density function

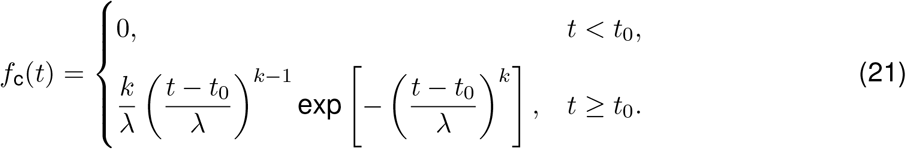

For the full population, the cumulative distribution function is

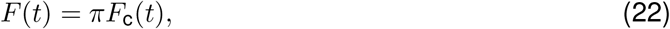

where *π* ∈ [0, 1] denotes the fraction of cells competent to undergo the transition.

For *π <* 1, *F* (*t*) is a defective distribution function with limiting value

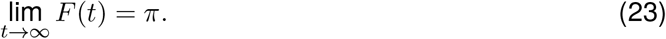

Accordingly, if the non-competent fraction 1−*π* is interpreted as never transitioning, then the unconditional transition-time distribution over the full population has point mass at infinity, and its mean and variance are not finite. For this reason, the moments derived below are those of the competent subpopulation, conditional on eventual transition.

Let

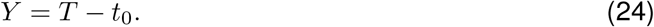

Then *Y* has the standard Weibull distribution

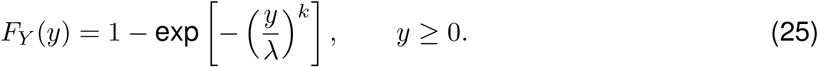

The *n*-th raw moment of *Y* is

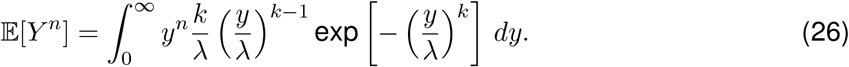

With the substitution

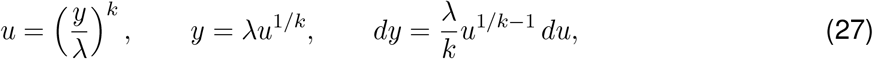

this becomes

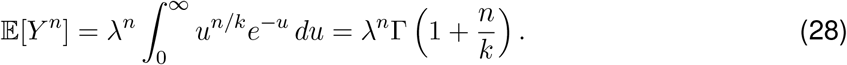

Therefore,

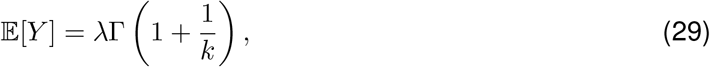

And

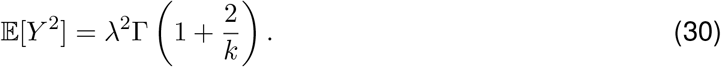

Since *T* = *t*_0_ + *Y*, the mean transition time among competent cells is

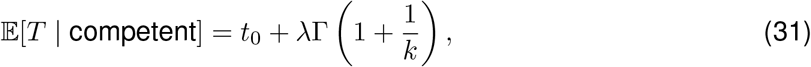

and the variance is

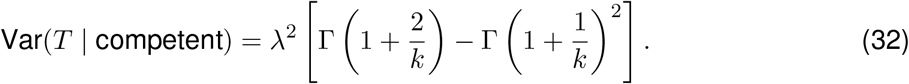

The conditional mean transition time is therefore

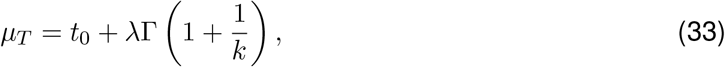

and the conditional variance is

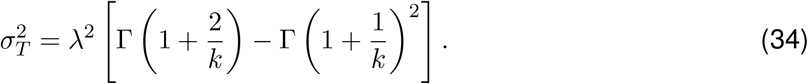

These expressions show that *t*_0_ enters additively into the mean and does not affect the variance, whereas *λ* scales both the mean and the variance. By contrast, the effect of *k* is nonlinear and is most naturally interpreted through the hazard function. The parameter *π* does not affect the conditional mean or variance among competent cells, but instead determines the asymptotic fraction of the population that undergoes transition.

